# The leader proteins of Theiler’s virus and Boone cardiovirus use a combination of Short Linear Motifs (SLiMs) to target RSK kinases to the nuclear pore complex

**DOI:** 10.1101/2025.04.09.647954

**Authors:** Belén Lizcano-Perret, Fanny Wavreil, Martin Veinstein, Camille Duflos, Romane Milcamps, Frédéric Sorgeloos, Thomas Michiels

## Abstract

Unrelated pathogens, including viruses and bacteria, use a common DDVF-like short linear motif (SLiM) to interact with cellular kinases of the RSK (p90 S6 ribosomal kinase) family. Such a “DDVF” SLiM occurs in the leader (L) protein encoded by picornaviruses of the genus *Cardiovirus*, including Theiler’s murine encephalomyelitis virus (TMEV), Boone cardiovirus (BCV), and Encephalomyocarditis virus (EMCV). The L-RSK complex is targeted to the nuclear pore, where RSK triggers FG-nucleoporins hyperphosphorylation, thereby causing nucleocytoplasmic trafficking disruption. In this work, we identified a second SLiM in the L proteins of TMEV and BCV, which enables the L-RSK complex to interact with RAE1 at the level of the nuclear pore complex. AlphaFold predictions suggest that the RAE1-interacting SLiM of L proteins is analogous to that found in unrelated viral proteins such as ORF6 of SARS-CoV-1/2, ORF10 of Kaposi sarcoma-associated herpes virus (KSHV), and the matrix (M) protein of vesicular stomatitis virus (VSV). Co-immunoprecipitations confirmed the interaction between BCV L and RAE1 and competition experiments revealed that L can compete with ORF6 for RAE1 binding, suggesting that BCV and TMEV L proteins interact with RAE1 via the same docking site as M, ORF6, or ORF10. This RAE1 binding SLiM tentatively named “M-acidic”, is predicted to occur in other viral proteins such as Rift valley fever virus NSs as well as in cell proteins such as NXF1. BCV and TMEV L proteins use a combination of two independent SLiMs to hijack cellular kinases and retarget those kinases toward the nuclear pore complex.

**Importance:** Protein-protein interactions are critical to regulate cell physiology. Short linear motifs (SLiMs) are unstructured protein sequences, which usually bind to structured domains of partner proteins. They typically mediate low affinity, transient interactions, which are particularly suitable for fine tuning cell physiology or helping cells to react promptly to stress situations. Owing to their fast replication and to the high error rate of their polymerases, viruses, particularly RNA viruses are prone to acquire SLiMs that mimic cellular SLiMs and thereby interfere with host cell signaling. In this work, we show that the leader (“L”) protein expressed by some cardioviruses (*Picornaviridae* family) uses two SLiMs in combination, which are individually shared by other pathogens: the first one, described previously, enables the L protein to hijack cellular kinases named RSKs, and the second one described in this work enables the L-RSK complex to target proteins RAE1 and NUP98 in the nuclear pore complex.

## Introduction

As obligate parasites, viruses developed a number of strategies to hijack cellular functions to their own benefit. Many such strategies are based on protein-protein interactions. Recent studies highlighted the importance of short linear motifs (SLiMs) in transient, low affinity protein-protein interactions that are instrumental for fine-tuning critical cellular signal transduction pathways (1). SLiMs are typically very short (≈ 3-10 amino acid-long) disordered protein motifs which trigger protein-protein interactions with structured domains of their protein partner (2, 3). Given the high error rate of their RNA-dependent RNA polymerases giving rise to quasi-species, RNA viruses are particularly prone to generate SLiMs in unstructured regions of their proteins, enabling them to take advantage of or to interfere with the regulation of cellular pathways, through SLiM mimicry (4, 5).

The genus *Cardiovirus*, in the *Picornaviridae* family encompasses human and animal viruses such as Saffold virus (SAFV), Encephalomyocarditis virus (EMCV), Theiler’s murine encephalomyelitis virus (TMEV or Theiler’s virus) and Boone cardiovirus (BCV). The leader (L) protein of these viruses is a very short protein (≈ 70-80 amino acids) cleaved-off from the amino-terminal end of the polyprotein encoded by these viruses. It acts as a multifunctional accessory protein (6). Theiler’s virus L protein was shown to be instrumental in the initiation of persistent infections of the central nervous system (7). In infected cells, L inhibits interferon (IFN) gene transcription, prevents PKR activation, and interferes with nucleocytoplasmic trafficking of host proteins (7-18). Moreover, the L protein of cardioviruses was recently shown to promote viral release in extracellular vesicles (19, 20).

PKR inhibition and nucleocytoplasmic traffic perturbation by L strongly depend on the ability of the L protein to interact with cellular kinases of the p90 S6 ribosomal kinase (RSK) family (21). Interaction of L with RSKs is mediated by a newly identified SLiM whose sequence is D/E-D/E-V-F, further referred to as “DDVF”, which has been shown to be shared by proteins from highly unrelated pathogens: L of cardioviruses (small non-enveloped RNA viruses), ORF45 and ORF11 of Kaposi sarcoma-associated herpes virus (KSHV) and Varicella-Zoster virus respectively (enveloped DNA viruses), and YopM of *Yersinia* species. The latter bacterial protein can contact RSK after being injected by the bacterium into eucaryotic cells through a type-III secretion system (22). The DDVF SLiM of L, ORF45, ORF11 and YopM was shown to bind RSKs through the same interface (21, 23). Since these pathogens’ proteins are not homologous, they likely acquired the SLiM by convergent evolution.

The L protein of TMEV induces a dramatic nucleocytoplasmic perturbation in infected cells. This activity was shown to depend on L-RSK interaction as it was abrogated upon either RSK depletion by CRISPR-Cas9 or F-to-A mutation of DDVF in L (L^F48A^), a mutation that prevents L interaction with RSK. Proximity labeling experiments showed that the L-RSK complex is targeted to the nuclear envelope, likely through interaction of L with components of the nuclear pore complex (NPC). L acts by redirecting part of the cellular RSKs toward unconventional substrates, such as phenylalanine-glycine nucleoporins (FG-NUPs), thereby triggering the aberrant diffusion of nuclear and cytoplasmic proteins between the two compartments (24). Interestingly, the M60V mutation in L was shown to prevent L-RSK complex targeting to the nuclear envelope, although this mutation did not affect the interaction between L and RSK (21).

In this work, we aimed to identify the determinants which target the L-RSK complex toward the NPC. We identified a second SLiM in the L protein, which also appears to be shared by other viral proteins and acts in combination with the DDVF SLiM to retarget RSK toward components of the NPC. This work illustrates that viruses can acquire independent SLiMs and use them in a combinatorial manner to promote their replication and to escape host immune responses.

## Results

### Theiler’s virus L protein maps to the nucleoplasm and the nuclear envelope

Previous proximity labeling data showed that the L protein fused to a biotin-ligase triggered the biotinylation of NPC components. This was not the case when L bore the M60V mutation, pointing to the idea that the L^M60V^ mutant no longer localized at the NPC (NPC)(24). To confirm the ability of L to localize at the nuclear envelope in a more direct setting, we used the split-GFP system (25). The sequence coding for the short GFP segment (16-amino acid “S11” segment) was cloned in the viral genome as an N-terminal extension of L. Viruses expressing S11-L^WT^ as well as mutants S11-L^M60V^ and S11-L^F48A^ were used to infect HeLa^S1-10^ cells expressing the large GFP segment “S1-10”. Results shown in Fig 1 confirm the ability of S11-L^WT^ and S11-L^F48A^ but not S11-L^M60V^ to be targeted to the nuclear envelope. The L^F48A^ mutant, which no longer interacts with RSK, showed clearer localization to the nuclear envelope compared to L^WT^. This clearer nuclear envelope localization likely results from the fact that L^F48A^ is less retained in the nucleoplasm than L^WT^ through interaction with RSK (Fig. 1).

**Figure 1.**
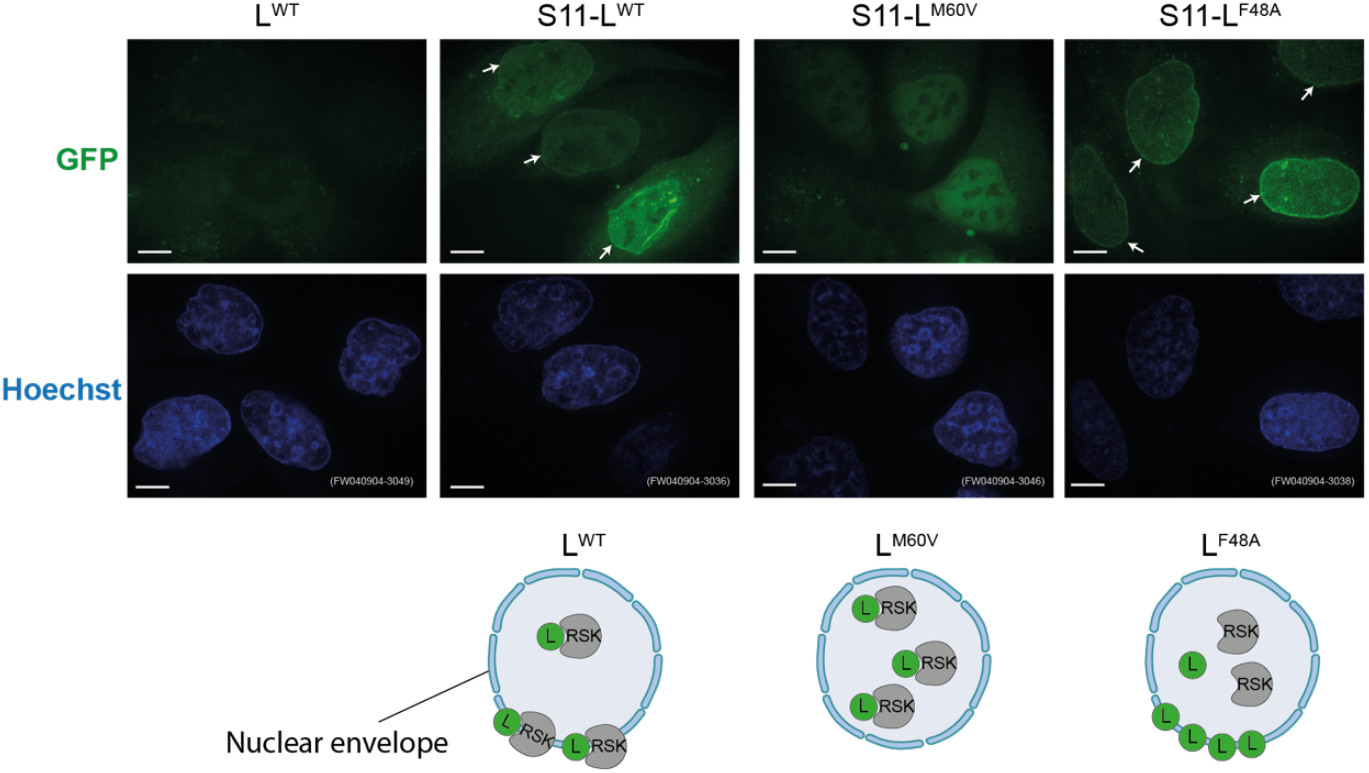
L^WT^ and L^F48A^, but not L^M60V^ localize at the nuclear envelope in infected cells. Confocal microscopy images of live HeLa^S1-10^ cells infected with S11-L viruses for 14h at an MOI of 5 PFU/cell. Arrows point at L protein localization at the nuclear envelope. Scale bar: 10µm

### Short linear motifs near the C-terminus of TMEV and BCV leader proteins are sufficient to target GFP to the NPC

Split GFP as well as full GFP fusions were used to map the nuclear envelope targeting motif in L proteins of TMEV, BCV and EMCV, which were chosen as evolutionarily distant cardioviruses. Therefore, pcDNA3 vectors expressing various portions of L proteins or mutants thereof fused to GFP or to the S11 GFP segment were transfected in HeLa cells or in HeLa^S1-10^ cells, respectively. Twenty-four to 36 hours post-transfection, cells were examined for green fluorescence at the level of the nuclear envelope. Results presented in Fig. 2A show that a nuclear envelope targeting motif lies in a 20 amino acid-long region located at the C-terminus of TMEV L (residues 57-76) and near the C-terminus of BCV L (residues 62-81). Of note, clear nuclear envelope localization was mostly visible in cells that expressed small amounts of the fusion protein. Transfected cells expressing higher protein fusion levels had fluorescence throughout the cell. In the case of EMCV, both N-and C-terminal fusions were tested with various parts of the protein and none of the fusions was readily targeted to the nuclear envelope, except the full-length protein. We conclude that the nuclear envelope targeting signal of EMCV is likely multipartite or conformational. In contrast, for TMEV and BCV, the targeting signal lies in a discrete motif predicted to be unstructured and thus might be considered as a SLiM. A highly similar sequence also occurs in the L protein of Saffold virus (SAFV), a human cardiovirus related to TMEV (Fig. 2B).

**Figure 2.**
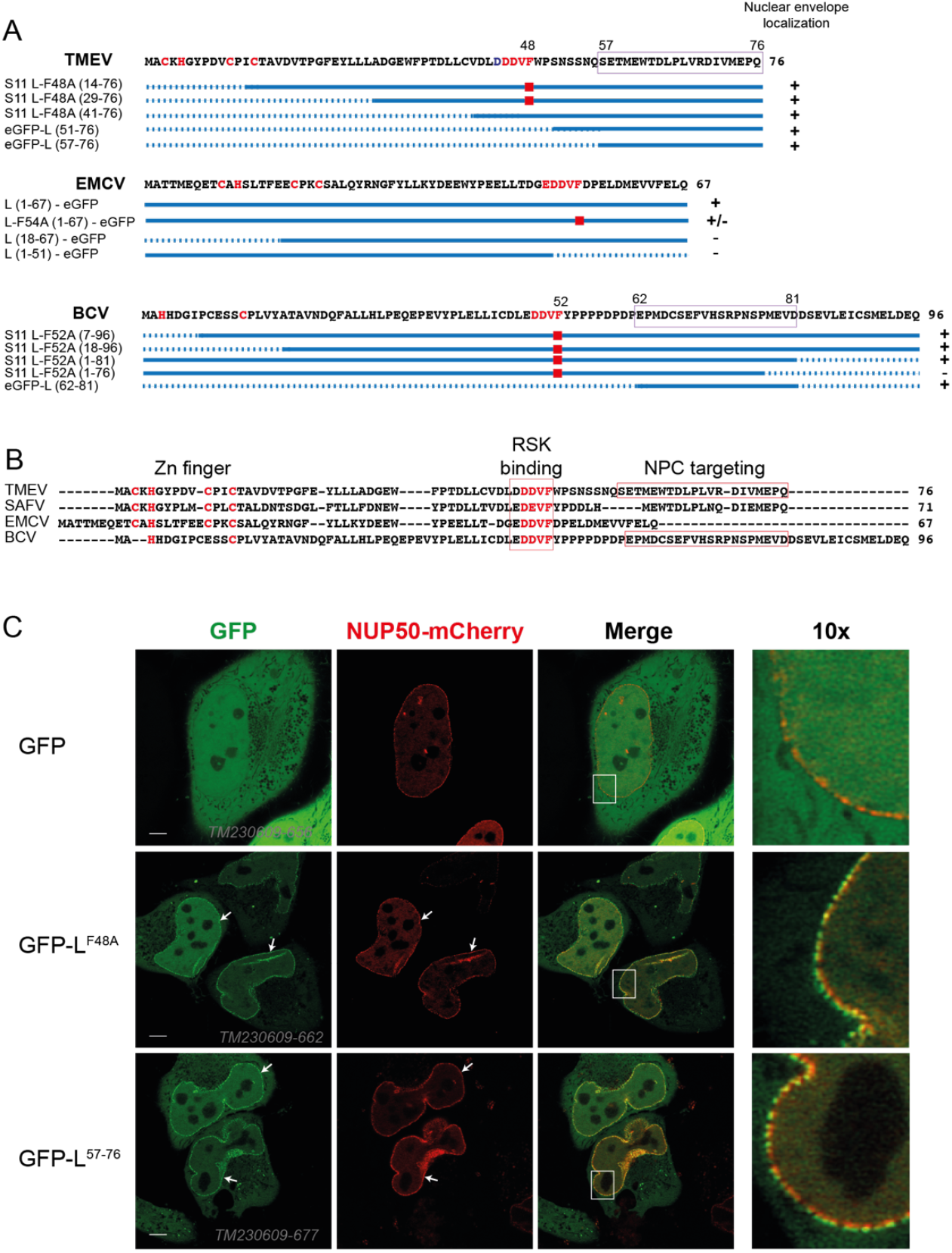
A 20 amino acid peptide of cardiovirus L is sufficient to target GFP to the NPC. **(A)** Mapping of the Cardiovirus L protein region sufficient for GFP-fusion localization to the NPC. Tested constructs are shown (solid line: region included in the constructs; dotted line: deleted region). Right of the constructs, + denotes nuclear envelope localization and -denotes absence of nuclear envelope localization. Red squares mark the F-to-A mutation in the RSK binding motif. A more detailed description of expression plasmids is presented in supplemental Table A1. (B) Alignment of representative cardiovirus L proteins with outline of the Zn finger, the DDVF-like SLiM involved in RSK binding, and the identified NPC localization SLiM (framed). SAFV: Saffold virus, a human virus related to TMEV. **(C)** High resolution confocal microscopy images of live HeLa cells co-transfected with constructs expressing NUP50-mCherry and GFP-L variants. Right panels show a 10-fold magnification of the white rectangle area shown in the merge image. Scale bar: 10µm

GFP-L fusions exhibited a dotted pattern at the level of the nuclear envelope, suggesting that L might interact with components of the NPC, as was also suggested by the proximity labelings described previously (24). We thus examined the co-localization of GFP-L and Nup50-mCherry in HeLa cells co-transfected with vectors expressing the two constructs. High-resolution microscopy revealed clear co-localization of GFP-L constructs with Nup50-mCherry (Fig. 2B), indicating that the leader protein motif was targeting GFP to the NPC. The 20 C-terminal amino acids of TMEV L were sufficient to target GFP to the NPC (GFP-L^57-76^ in Fig. 2B).

### TMEV L NPC targeting correlates with protein toxicity

To delineate the NPC-targeting motif at higher resolution, we performed Alanine-scan mutagenesis of the TMEV L^57-76^ peptide fused to GFP (Fig. 3A). Cells transfected with the various mutants were examined by fluorescence microscopy and co-localization with mCherry-Lamin was quantified by confocal microscopy in cells that expressed low green fluorescence levels. As shown in Fig. 3A, residues that strongly affected NPC localization of the fusion protein were M60, E61, W62, T63, P66, D70 and M73, and to a lesser extent, V68. Except for the latter, these residues remarkably correlate with those previously selected after random L mutagenesis, as residues involved in L protein toxicity (Fig. 3B) (26). These data strongly suggest that L protein toxicity is linked to the ability of L to target the NPC.

**Figure 3.**
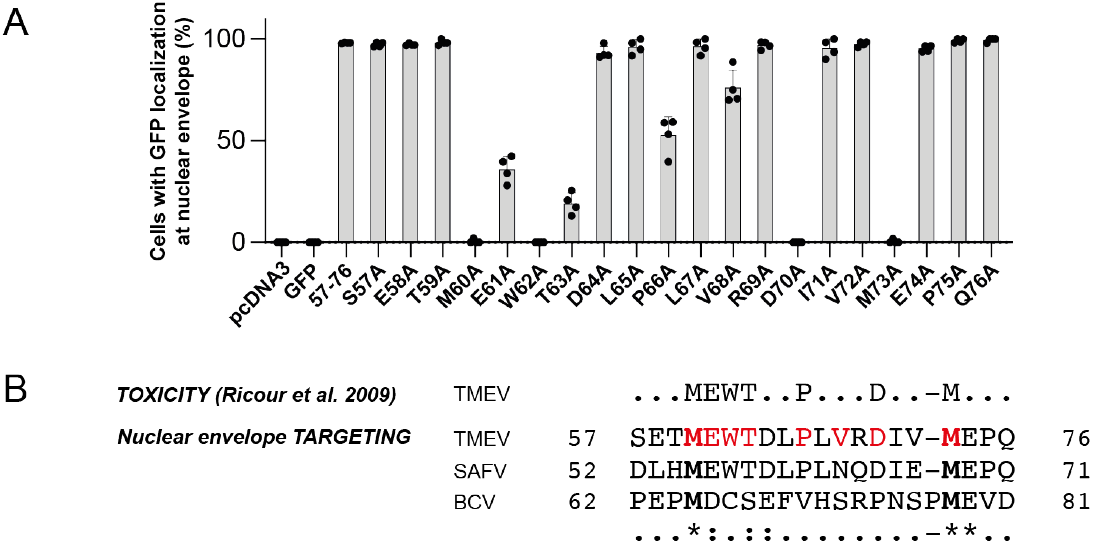
TMEV L protein localization at the NPC correlates with its cellular toxicity. **(A)** Alanine-scan mutagenesis of GFP-L^57-76^. Graph showing the percentage of cells with GFP localization at the nuclear envelope (mCherry-Lamin co-localization), n = 4. **(B)** Sequence alignment showing key amino acids required for TMEV L toxicity, TMEV and BCV L NPC localization, and the corresponding peptide in SAFV L. Red residues are amino acids important for NPC localization.

### Mutation of methionines in the NPC-targeting motif suppresses the ability of TMEV, SAFV and BCV L proteins to perturb nucleocytoplasmic trafficking

The critical amino acids for GFP-L^57-76^ localization at the NPC were M60, E61, W62, T63, P66, D70 and M73, and V68 (Fig. 3A). Among these residues, two methionines (M60 and M73 for TMEV) are strictly conserved in cardiovirus L proteins (Fig. 3B). To assess whether the conserved methionines contained in the NPC targeting motif are critical for nucleocytoplasmic perturbation during viral infection, TMEV was engineered to express the L protein of SAFV or that of BCV. Corresponding mutant viruses containing mutations in the conserved methionines, which are important for NPC localization, were also constructed. In the case of TMEV and SAFV, only the first M residue was mutated since the second M residue is likely critical for viral polyprotein cleavage. HeLa cells expressing GFP-NES and RFP-NLS were infected with those viruses and followed by live cell imaging to document the impact of infection on nucleocytoplasmic trafficking (Fig. 4). As expected, mutation of M60 in TMEV L, of M55 in SAFV L, and of M64 and M78 in BCV L strongly affected the ability of L to disrupt nucleocytoplasmic trafficking. Interestingly, for BCV L, on the one hand, redistribution of GFP-NES and RFP-NLS occurred as early as 4h30 post-infection compared to ∼6h30 for TMEV L, and on the other hand, mutation of a M64 or M78 alone was not sufficient to abrogate L-mediated nucleocytoplasmic trafficking perturbation. This suggests a higher affinity for BCV L than TMEV L for the NPC.

**Figure 4.**
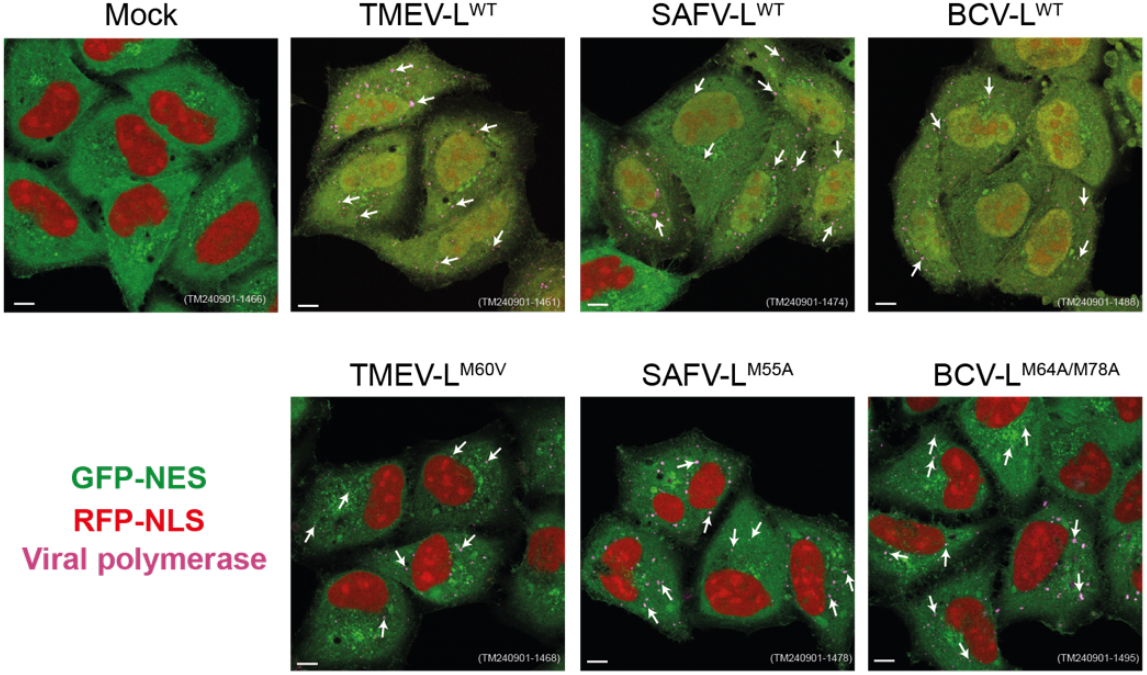
Conserved methionines are important for nucleocytoplasmic traffic disruption. Confocal microscopy of HeLa cells expressing GFP-NES and RFP-NLS infected with TMEV derivatives expressing TMEV L, SAFV L or BCV L for 12 h at an MOI of 5 PFU/cell. Cells were fixed and viral polymerase (3D) was immunolabeled as control of infection (purple dots + arrows). Scale bar: 10µm.

### Predictions suggest that L proteins interact with NPC components through a “M-acidic” SLiM shared by other pathogens’ proteins

It was reported that unrelated viral proteins, such as the matrix (M) protein of vesicular stomatitis virus (VSV) (27, 28), the ORF10 protein of KSHV (29, 30) and the ORF6 protein of SARS-CoV-1 and SARS-CoV-2 (31-34) interact with the complex formed by nucleoporins “ribonucleic acid export 1” (RAE1) and “nucleoporin 98” (NUP98) at the NPC. In all these proteins, a methionine surrounded by acidic residues and sometimes preceded by a proline (Fig. 5A) was shown to critically contact a hydrophobic pocket formed at the interface of RAE1 and NUP98. This SLiM is therefore further referred to as the “M-acidic” SLiM. This hydrophobic pocket is thought to be RAE1’s interface for binding and exporting RNAs via their phosphate backbones. Thus, these pathogens’ proteins use SLiM mimicry to bind RAE1 and compete for RNA binding by mimicking RNA phosphates with acidic residues, thereby blocking mRNA export (27, 33). Interestingly, the key methionines needed for L protein localization to the NPC are contained in motifs resembling the RAE1-interacting M-acidic SLiM of SARS-CoV ORF6 and of the other pathogens’ proteins (Fig 5A). Of note, like those proteins, cardiovirus L proteins have also been shown to inhibit mRNA export (14). We thus used AlphaFold 2 multimer (35) to predict whether either full-length L proteins or the 20-mer peptides (Fig 5B-C) of TMEV and BCV interact with RAE1/NUP98 through a similar interface as the other pathogens’ proteins. Interestingly, BCV L was predicted to interact with RAE1/NUP98 through either the M64 or M78 residue and TMEV L mostly through the M60 residue (Fig 5B-D). The predicted interfaces are similar to those of SARS-CoV ORF6, KSHV ORF10 and VSV M, with the methionine residue inserted in a hydrophobic pocket of RAE1 (Fig 5C).

**Fig. 5.**
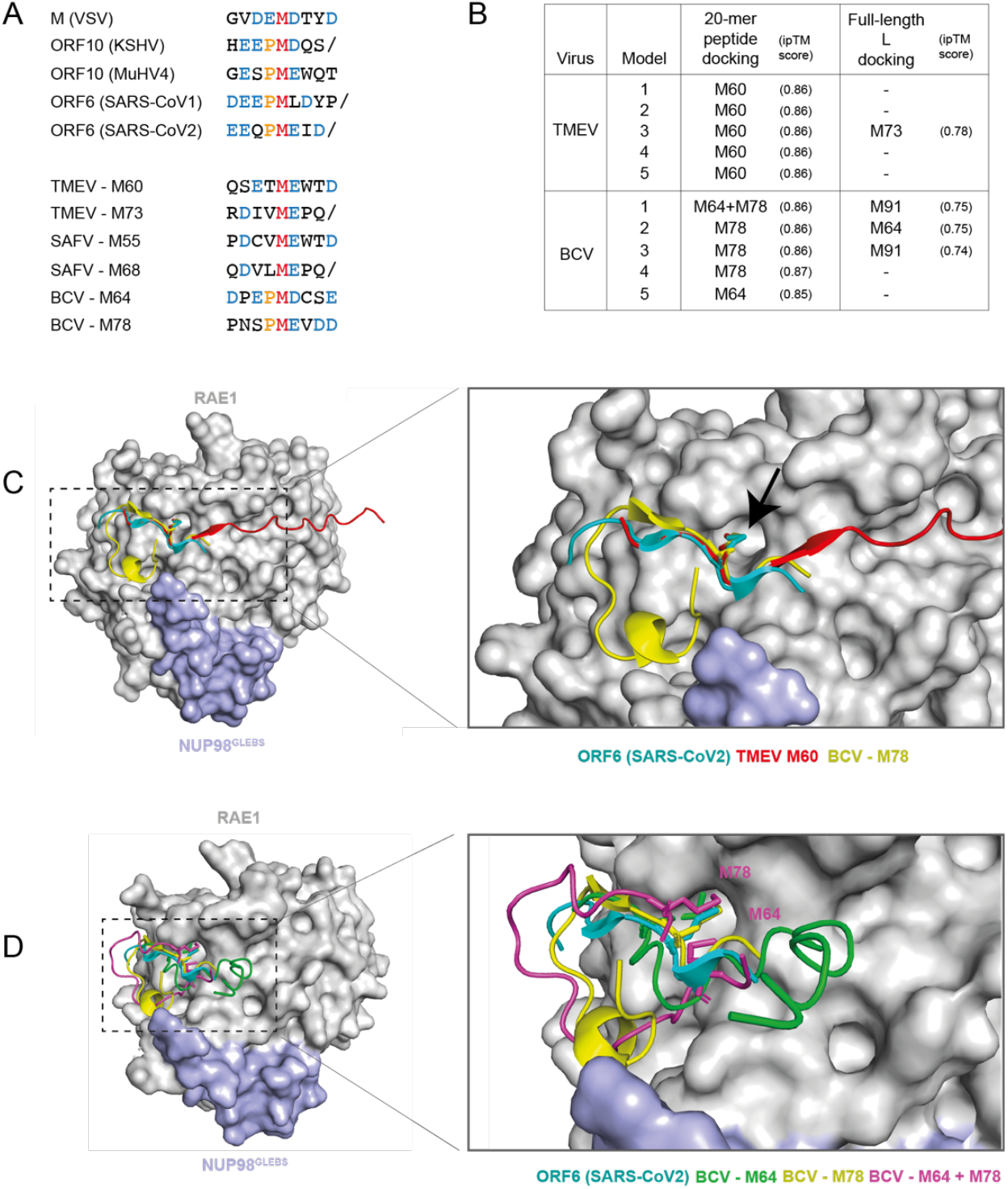
Structural predictions indicate that cardiovirus L proteins could interact with RAE1 and NUP98 through a similar interface as ORF6 from SARS-CoV2. (A) Sequence alignment of protein motifs predicted to or shown to bind RAE1/NUP98. Sequences are from M (VSV), ORF10 (KSHV or MuHV4), ORF6 (SARS-CoV1/2) and L (TMEV, SAFV and BCV). (B) AlphaFold 2 multimer predictions of interfaces between TMEV and BCV full-length L or 20-mer peptides with RAE1 and NUP98^GLEBS^ regions used for crystallography of the RAE1-NUP98-ORF6 complex (PDB: 7VPH). Interface predicted template modelling scores (ipTM) > 0.8 are considered accurate high-quality predictions. (C) Structural alignment of SARS-CoV2 C-terminal tail with RAE1 and NUP98^GLEBS^ (PDB: 7VPH) and the AlphaFold predictions of the L 20-mer peptides of TMEV and BCV, showing similar insertion of the methionine (black arrow) in the hydrophobic pocket of RAE1. (D) multiple BCV models showing either M64 or M78 insertion in the hydrophobic pocket.

### L of BCV and TMEV compete with ORF6 to interact with RAE1

To assess the ability of TMEV and BCV L proteins to interact with RAE1 and NUP98, co-immunoprecipitations were performed from 293T cells transfected with constructs expressing FLAG-L (TMEV or BCV) or FLAG-ORF6 (SARS-CoV2) taken as a positive control. Results presented in Fig 6A show that endogenous RAE1 readily co-immunoprecipitated with BCV L but hardly with the mutant BCV L^M64A/M78A^, in line with the data showing the importance of M64 and M78 for nucleocytoplasmic traffic perturbation (Fig 4). NUP98 co-immunoprecipitation was unclear though suggested in some experiments. In the case of TMEV L, no interaction was detected by co-IP. Our interpretation is that L of BCV has a higher affinity for RAE1 than TMEV L, as suggested by the more pronounced nucleocytoplasmic trafficking perturbation induced by L of BCV, i.e. double methionine mutation needed to fully inhibit the effect.

**Figure 6.**
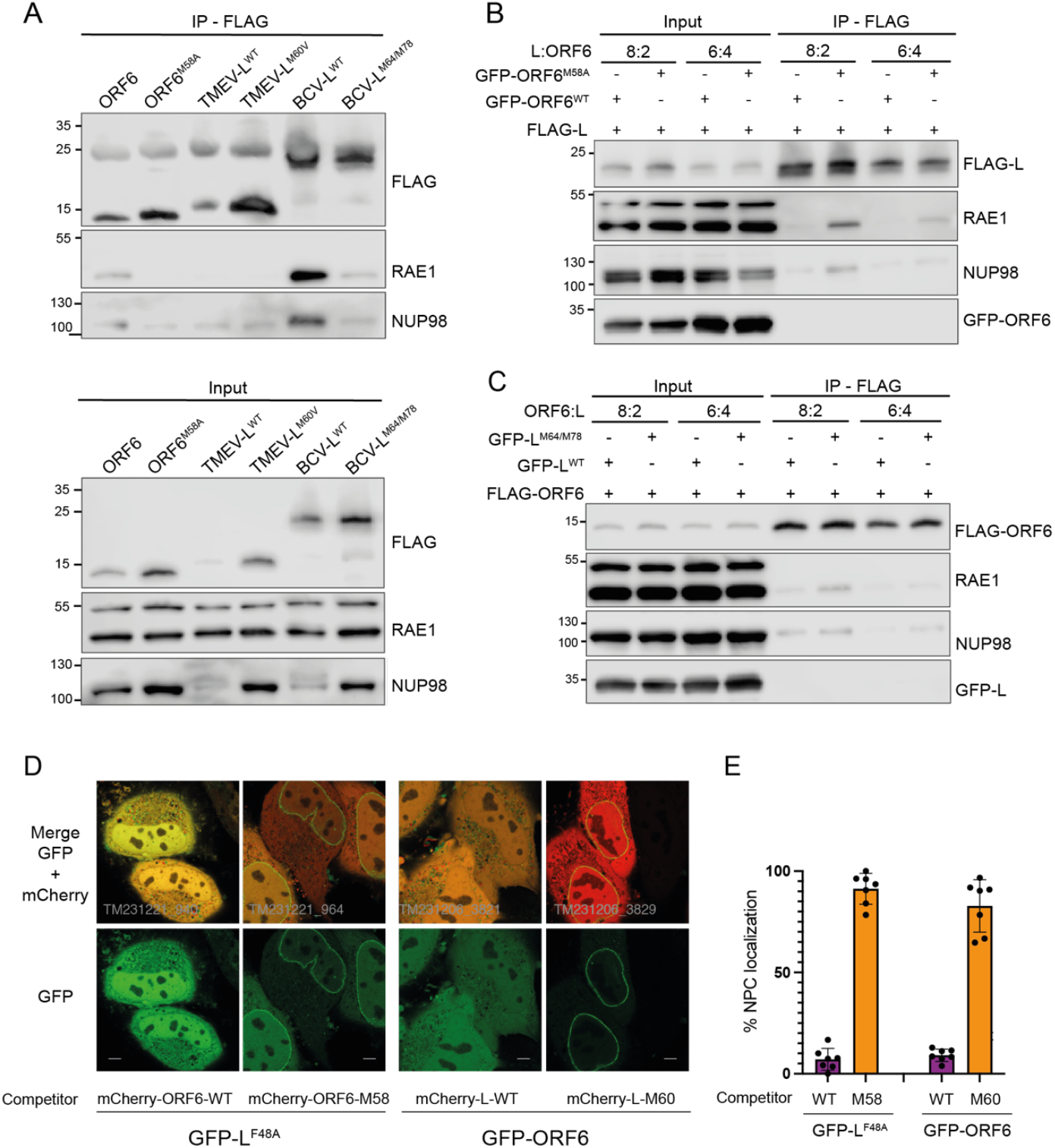
TMEV and BCV L compete with SARS-CoV-2 ORF6 for RAE1 interaction. **(A)** Western blots showing the detection of endogenous RAE1 and NUP98 after immunoprecipitation of FLAG (FLAG-L or FLAG-ORF6) from 293T cells transfected for 24h with plasmids expressing the indicated FLAG-tagged construct. **(B-C)** Western blots showing the competition for RAE1 co-immunoprecipitation, between BCV L and SARS-CoV-2 ORF6. 293T cells were co-transfected for 48h with constructs expressing BCV FLAG-L and GFP fused to the 20-mer NPC targeting peptide of SARS-CoV-2 ORF6 (ORF6^42-61^) (B) or with constructs expressing FLAG-ORF6 and GFP fused to the BCV L^62-81^ NPC targeting peptide (C). FLAG-tagged constructs and competitors were transfected at different ratios: 8:2 and 6:4. Competitors contained either the WT or the indicated mutant NPC targeting peptide. **(D)** High-resolution microscopy images showing examples of cells displaying clear competition for NPC targeting, between SARS-CoV-2 ORF6 and TMEV L. Left four panels: HeLa cells were co-transfected for 36h with 25 ng of plasmid expressing GFP-L^F48A^ and 125 ng of plasmid expressing mCherry-ORF6^42-61^ either wild type (WT) or carrying the M58A mutation. Right four panels: HeLa cells were co-transfected for 36h with 25 ng of plasmid expressing GFP-ORF6 and 125 ng of plasmid expressing mCherry-L^57-76^ either wild type (WT) or carrying the M60V mutation. (E) Quantification of microscopy images showing the percentage of mCherry-positive cells which displayed distinct NPC localization of GFP. Counted on 10 high-resolution images per transfected well, from a total of 6 or 7 transfected wells.

We next tested whether some competition exists between BCV L and SARS-CoV-2 ORF6 for RAE1 binding in co-immunoprecipitation experiments. A construct expressing FLAG-L of BCV was transfected in 293T cells along with vectors expressing GFP fused to the 20 C-terminal residues of ORF6, which have been shown to be sufficient for targeting the fusion protein to the nuclear envelope. FLAG-L was co-expressed with GFP-ORF6^WT^ or GFP-ORF6^M58A^ mutant, which no longer interacts with RAE1. After FLAG-L immunoprecipitation, RAE1 co-immunoprecipitated in higher amounts from cells expressing the mutant GFP-ORF6^M58A^ than from cells expressing wild type GFP-ORF6 (Fig. 6B). Conversely, co-expression of GFP fused to the NPC-targeting 20-mer of BCV L decreased RAE1 co-immunoprecipitation with FLAG-ORF6 more than the GFP-L^M64A/M78A^ mutant construct (Fig. 6C). These data show that BCV L and ORF6 compete to bind RAE1.

RAE1 co-immunoprecipitation with TMEV L was not detectable, likely because affinity of TMEV L for RAE1 is lower than that of BCV L. Therefore, we could not use the above strategy to assess whether TMEV L competes with ORF6 for NPC binding. We thus used immunofluorescent microscopy to test whether competition for NPC localization exists between TMEV L and SARS-CoV-2 ORF6. Since wild type L and, to a lesser extent, ORF6 are toxic and affect mRNA translation (14, 31), we used the less toxic mutants or truncated proteins to perform the competition experiments. Thus, a construct expressing GFP-L^F48A^ was transfected in HeLa cells with constructs expressing mCherry fused to the C-terminal 20 amino acids of ORF6, which are sufficient for NPC targeting. The C-terminus of ORF6 was either WT or carrying the M58A mutation that was shown to abrogate NPC targeting (34). Competitions were also performed the other way around, using vectors that express GFP-ORF6 and fusions between mCherry and the C-terminal peptide of TMEV L (L57-76), which was either WT or carrying the M60V mutation. Although these experiments have clear limitations due to the greater impact of wild type than mutant competitors on GFP expression, results shown in Fig 6D-E were reproducible and strongly suggested that TMEV L also competes with ORF6 for NPC targeting, via a M-acidic SLiM binding to RAE1.

### Different impact of L, ORF6, and ORF10 on nucleocytoplasmic traffic

Our results indicate that BCV L and likely TMEV protein interact with RAE1 and NUP98 via a M-acidic SLiM as other pathogen’s proteins, namely the M protein (VSV), ORF6 (SARS-CoV1/2) and ORF10 (KSHV). All these pathogen’s proteins, including cardiovirus L, have been shown to inhibit mRNA export (14, 30, 31, 33, 36, 37). For some of these proteins not only RNA export is inhibited but the localization of nuclear or cytoplasmic proteins is also disturbed. SARS-CoV2 ORF6 was shown to block STATs nuclear import (34), VSV M protein was shown to induce the cytoplasmic diffusion of hnRNPA1, hnRNPK, and hnRNPC1/C2 (38), and cardiovirus L protein was shown to induce a redistribution of the polypyrimidine tract-binding protein (PTB), the interferon regulatory factor 3 (IRF3) as well as of a GFP-NLS fusion protein (12). Even though all these pathogen’s proteins interact with RAE1 and NUP98 via the same SLiM, only the L protein possesses the DDVF-SLiM allowing RSK recruitment and FG-NUPs phosphorylation. Therefore, we wondered whether these different proteins had a similar impact on nucleocytoplasmic trafficking. We thus transfected FLAG-L (TMEV and BCV), FLAG-ORF6 (SARS-CoV-2) and FLAG-ORF10 (KSHV) into HeLa cells expressing GFP-NES and RFP-NLS (Fig 7). Results show that only cardiovirus L proteins can disturb the localization of both GFP-NES and RFP-NLS. Although ORF6 and ORF10 were well expressed, they did not disrupt the localization of the fluorescent proteins. These results indicate that, even though these proteins interact with RAE1 and NUP98 via the same M-acidic SLiM, the impact of this on nucleocytoplasmic traffic is more dramatic for L proteins, which take advantage of the enzymatic activity of RSK.

**Figure 7.**
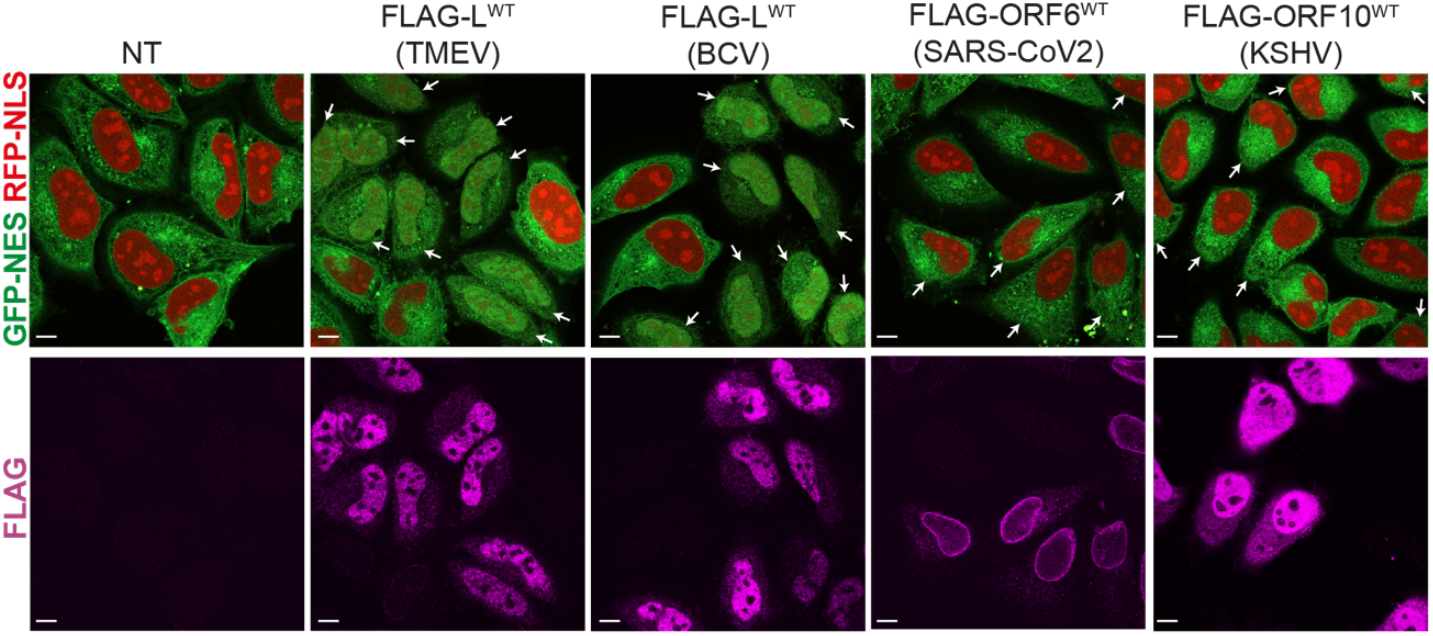
Cardiovirus L proteins disturb nucleocytoplasmic traffic to a bigger extent than ORF6 (SARS-CoV-2) and ORF10 (KSHV). Confocal microscopy of HeLa cells expressing GFP-NES and RFP-NLS transfected with FLAG-L (TMEV or BCV), FLAG-ORF6 and FLAG-ORF10. 24h post transfection, cells were stained with an anti-FLAG antibody. Arrows point at cells expressing FLAG tagged proteins. Scale bar: 10µm.

## Discussion

Our data identify a “M-acidic” SLiM located at or close to the C-terminus of Theiler’s virus and Boone cardiovirus L proteins, which mediates the interaction with components of the NPC, including RAE1. In infected HeLa cells, the leader protein of EMCV was shown to map around the nuclear envelope (39). Accordingly, EMCV L targeted a C-terminally fused GFP to the nuclear envelope in our experiments. We were however unable to delineate a short NPC targeting sequence in EMCV L, suggesting that the targeting signal is conformational or that interaction with the NPC involves more than one segment of the EMCV L protein.

In the motif identified for TMEV and BCV L proteins, conserved methionine residues are essential for NPC targeting. Other viral proteins were previously shown to target the RAE1-NUP98 complex, including the matrix (M) protein of VSV, the ORF10 protein of KSHV, and the ORF6 protein of SARS-Co-V. Strikingly all these viral proteins were shown to dock in the RAE1-NUP98 complex through a very similar interface, which involves the insertion of a methionine residue in a hydrophobic pocket of RAE1 (27, 29, 33). AlphaFold predictions and competition experiment results (see Fig 6) suggest a similar interaction of L with RAE1 through the M-acidic SLiM. It is however unclear whether the two methionines found in cardiovirus L proteins, which are both important for binding, alternatively insert in the same pocket or contact distinct RAE1-NUP98 complexes. It is also possible that one of the methionines is tightening the interaction by binding elsewhere on the surface of RAE1-NUP98.

SLiM-mediated interaction of unrelated viral proteins with the same NPC component provides a typical example of convergent evolution. Given their high burst rate viruses can easily evolve to include SLiMs in their proteins, in particular RNA viruses which use error-prone polymerases.

Viral SLiMs typically evolve to mimic SLiMs which mediate interaction between cellular proteins. Such SLiM mimicry by pathogens has been recently documented for the DDVF SLiM, which was found to mimic a SLiM occurring in proteins regulating the MAP kinase pathway enabling them to interact with RSK kinases (40). Interestingly, the conserved SLiM-docking site at the surface of RAE1-NUP98 was shown to be a site where mRNA binds during their export to the cytosol. In this case, M-acidic SLiMs are thus mimicking RNAs instead of short protein motifs. Nevertheless, we used AlphaFold to predict whether similar M-acidic SLiMs docking in RAE1-NUP98 might occur in human proteins or in additional viral proteins that were reported to interact with both RAE1 and NUP98, in the IntAct and BioGRID databases (41, 42). Since longer protein sequences significantly increase prediction time and reduce protein structure prediction accuracy (43), we excluded proteins with lengths exceeding 1000 residues, reducing the dataset to 62 proteins. For these 62 proteins, we conducted AlphaFold2 multimer v3 predictions using their sequences alongside those of RAE1 and NUP98, as derived from PDB 7VPH. Two proteins were predicted to interact with RAE1-NUP98 through a M-acidic SLiM: Rift valley fever virus (RVFV) protein NSs (residue M250 in sequence DVE<u>M</u>ESEEE) and the human protein Nuclear RNA export factor 1 (NXF1/TAP) (residue M61 in the sequence GDVA<u>M</u>SDAQD).

RVFV NSs interaction with RAE1-NUP98 was detected by high throughput tandem affinity purification (44) but efforts to individually co-immunoprecipitate RAE1 with NSs were reportedly unsuccessful (45), likely because the affinity of the SLiM of NSs for RAE1 is under the threshold for successful co-immunoprecipitation, as was the case for TMEV L in this work. The case of Nuclear RNA export factor 1 (NXF1/TAP) is interesting because this factor is reported to contribute to mRNA export by binding several partners in the NPC. It binds to FG-nucleoporins through its C-terminal ubiquitin-associated fold (UBA) as well as through its central region when forming a heterodimer with protein p15/NXT. In addition, NXF1 can also interact with RAE1 and binding to RAE1 was shown to be abrogated after deletion of the N-terminal 60 amino acids of NXF1 (46). In addition, M61 maps at the beginning of a region identified as the nuclear localization signal. Competition between viral proteins ORF6, ORF10, M, NSs and L with NXF1 for RAE1 binding would be in line with the observation that binding of these viral proteins to the NPC affect mRNA export (29, 31, 36, 47).

Although the L proteins of TMEV and BCV target the same NPC components as SARS-CoV ORF6 or KSHV ORF10, they were much more potent at disrupting the nucleocytoplasmic traffic. Indeed, in contrast to the other proteins that are believed to act by competition or by disturbance of NPC organization, L proteins make use of an enzymatic activity by recruiting RSK kinases, which in turn phosphorylate NPC components (24).

L proteins thus use a combination of 2 SLiMs, both being shared by other pathogens, to retarget RSKs toward the NPC, simultaneously affecting the MAP kinase pathway. It is interesting to note that such a combinatorial use of SLiMs is a potent and evolutionary simple way to hijack essential cellular processes.

## Material and Methods

### Cells

Cells referred to as HeLa cells in this work are HeLa M kindly provided by R.H Silvermann (48). The HeLa-LVX cells expressing RFP-NLS and GFP-NES were obtained as described previously (24). The HeLa^S1-10^ cells were obtained by transduction of HeLa M cells with the lentivirus BLP2 (see below), coding for the segment S1-10 of GFP. After transduction, a cellular clone having low amount of fluorescence in normal conditions, and high amount of green fluorescence upon addition of the S11 fragment was selected. 293T cells (49) used in this work (lentivirus production) were kindly given by F. Tangy (Pasteur Institute, Paris). HeLa and 293T cells used in this work were maintained in Dulbecco’s Modified Eagle Medium (DMEM high-glucose -Biowest) supplemented with 10% fetal calf serum (FBS, Hyclone-Cytiva), penicillin (100U/ml) and streptomycin (100μg/ml) (Gibco). BHK-21 cells used for virus production were maintained in Glasgow’s Modified Eagle Medium (GMEM, Gibco) with 2.6g/L of tryptose phosphate broth (Gibco), 10% newborn calf serum (Gibco) and penicillin (100U/ml)/streptomycin (100μg/ml).

All cell lines were cultured at 37°C, 5% CO_2._

### Plasmids, lentiviral vectors and viruses

Sequences referred to in this work as to GFP correspond to enhanced GFP (or EGFP). Sequences for the Split-GFP system (25) were PCR-amplified from plasmids kindly provided by Hilde van der Schaar (Univ. Utrecht). Plasmid mCherry-NUP50-N-10 and mCherry-LaminA-C-18 were gifts from Michael Davidson (Addgene plasmids # 55111 and # 55068). ORF10-cFLAG plasmid (30), kindly provided by Carissa Pardamean and Ting-Ting Wu (UCLA) served as a source of ORF10 DNA. Plasmid pCAD03 was obtained by cloning the ORF coding a catalytically inactive (GDD->DAA mutant) 3D polymerase of TMEV (strain DA1) in plasmid pOPTH. Expression plasmids were derived from pcDNA3 (Invitrogen). The lentiviral construct pLVX-EF1alpha-2xGFP:NES-IRES-2xRFP:NLS was a gift from Fred Gage (50)(Addgene plasmid # 71396). Lentiviral vector BLP2 was obtained by cloning the region expressing GFP S1-10 in pTM943 (51), a vector derived from pCCLsin.PPT.hPGK.GFP.pre. (52).

Theiler’s murine encephalomyelitis viruses were derived from KJ6, a variant of strain DA1 (accession JX443418.1) carrying capsid mutations selected to optimize L929 cell infection (53). Plasmids used in this work were constructed by standard methods and are compiled in supplemental Table A1.

### Lentivirus production and cell transduction

For lentivirus production, 293T cells were seeded in a 6-cm Petri dish. At ∼80% confluency cells were co-transfected with 5µg of lentiviral vector (pBLP2), 2.5µg pMDLg/RRE (Gag-Pol), 1.5µg of pMD2-VSV-G, and 1.25µg of pRSV-Rev using TransIT-LT1 (Mirus Bio). 24h and 48h post transfection, supernatant was collected and filtered (0.45µm). For transduction, 5,000 HeLa M cells were seeded in a 24-well plate and were transduced twice with 100µl of BLP2.

### Virus production and titration

TMEV derivatives used in this work were produced by reverse genetics using plasmids containing the full viral cDNA sequence. BHK-21 cells were electroporated with viral RNA (*in vitro* transcribed – RiboMax Promega P1300) using a Gene pulser apparatus (Bio-Rad) (1500V, 25µF, no shunt resistance). Supernatants were collected when cytopathic effect was complete (∼48h post-electroporation). Two to three freeze-thaw cycles were made to increase viral release from cells before clarifying the supernatants by centrifugation (20min, 1258xg). Viruses were stored at -80°C and titrated by plaque assay in BHK-21 cells as described (54).

### Co-Immunoprecipitations (Co-IP) and Co-IP competitions

293T cells were seeded in 6-cm dish as of 1,250,000 cells. 24h after the seeding, cells were transfected with 10µg of DNA using Mirus TransIT 2020 (Mirus Bio). For Co-IP competitions, 10µg of DNA were also used but at different ratios (8:2µg or 6:4µg). 40h post-transfection, cells were lysed for 30min on ice with 300µl of RIPA buffer (ref 23T1.2 -Roth) containing phosphatase/protease inhibitor (1 tablet for 10ml buffer, Pierce-ThermoScientific). Lysates were transferred to a tube, and 300µl of Tris-HCl 50mM pH8 was added to dilute the lysis buffer. Lysates where homogenized by passages through 21G needles and cleared by centrifugation at 12,000xg, 4°C for 10min. Supernatants were then incubated for 30min at 4°C with 25µl of A/G magnetic beads (Pierce) to remove unspecific binding. Supernatants were transferred to a new tube and a sample of 30µl per condition was mixed with 15µl of 3x Laemmli buffer (cell lysate control). The rest of the supernatant was incubated with anti-FLAG magnetic beads (M8823 – Sigma) for 2h at 4°C. Beads were washed three times with washing buffer (Mix 1:1 of RIPA Buffer 23T1.2 Roth and Tris-HCl 50mM pH8), resuspended in 35µl of Laemmli 1x and heated for 10min at 100°C to allow protein’s separation from the beads.

Supernatants were then conserved at -20°C.

### Western blot

Proteins in Laemmli buffer were heated at 100°C for 5minutes. Protein samples were then run in 12% Tricine SDS polyacrylamide gels (for FLAG-L or FLAG-ORF6 detection), 8% glycine SDS polyacrylamide gels (for NUP98 detection) or in 10% glycine SDS polyacrylamide gels. Proteins were then transferred to PVDF or nitrocellulose (only for FLAG-L or FLAG-ORF6) membranes. For FLAG-L and FLAG-ORF6 detection, membranes were treated with a signal enhancer (ref 46640 - ThermoScientific) prior to chemiluminescence detection. Membranes were blocked with TBS-0.1% Tween 20-5% milk (Regilait) for 1h at RT. Membranes were then incubated over-night with primary antibodies at proper dilution in TBS-0.1% Tween 20-5% milk. Primary antibodies used: anti-RAE1 (Ab124783, rabbit, 1/1000), anti-FLAG (F1804 Sigma, mouse, 1/5000) or anti-FLAG-HRP (A8592 Sigma, mouse, 1/2000), anti-NUP98 (N1038 Sigma, rat, 1/1000) and anti-GFP (66002-1-IG Proteintech, mouse, 1/10000). Membranes were washed 3 times with TBS-0.1% Tween 20 for 15min before incubation with secondary antibodies in TBS-0.1% Tween 20-5% milk for 1h at RT. Secondary antibodies used: HRP-conjugated anti-rabbit (Dako P0448 - 1/5000), HRP-conjugated anti-mouse (Dako P0447 - 1/5000), and HRP-conjugated anti-rat (CST 7077 - 1/5000). Membranes were washed 3 times with TBS-0.1% Tween 20, then once with TBS and revealed with SuperSignal West chemiluminescence substrate (Pico or Dura, ThermoScientific) or with Westar Supernova (Cyanagen). Images were taken with a cooled CCD camera (Odyssey FC- LiCor).

### S11/GFP-L plasmids transfection

HeLa M cells were seeded in 96-well plates (Greiner, 655866 Screenstar) at a density of 3,000 cells per well. 24h after seeding, cells were transfected with either 100ng DNA (50ng L-coding plasmid and 50ng of NUP50-mCherry coding plasmid) or 150ng DNA (50ng of L-coding plasmid and 100ng pcDNA3 or 1.5 ng of L-coding plasmid, 73.5ng of pcDNA3 and 75ng of Lamin-mCherry coding plasmid). 24h to 36h post-transfection, pictures of transfected cells were taken with either a spinning disk confocal microscope or a high-resolution LSM980 multiphoton microscope.

### Immunostaining

Cells seeded in a 96-well plate (Greiner, 655866 Screenstar) were fixed with PBS-4% PFA for 5min at RT before being permeabilized with PBS-0.2% Triton (ICN Biomedicals Inc.) for 5min. Cells were blocked with TNB blocking reagent (Perkin Elmer) for 1h at RT. Cells were then washed three times with PBS-0.1% Tween 20 before being incubated with primary antibody diluted in TNB for 1h at RT. Primary antibodies: anti-FLAG (F1804 Sigma, mouse - 1/800) and anti-3D-viral polymerase polyclonal rabbit antibody (See below - 1/1000). Cells were then washed three times with PBS-0.1% Tween 20 before being incubated with secondary antibody diluted in TNB for 1h at RT. Secondary antibodies: anti-mouse-Alexa Fluor-647 (A32728 - goat, 1/800) and anti-rabbit-Alexa Fluor-647 (A21245 - goat, 1/800). Cells were then washed three times with PBS-0.1% Tween 20 and kept in PBS-0.02% sodium azide.

### L-ORF6 competitions - Immunofluorescence

HeLa M cells were seeded in 96-well plates (Greiner, 655866 Screenstar) at a density of 3,000 cells per well. 24h after seeding, cells were transfected with 150 ng of DNA per well (25 ng of the GFP-L or GFP-ORF6 construct and 125 ng of the mCherry-L or mCherry-ORF6 competitor). High-resolution images were taken with a LSM980 Zeiss microscope with the following settings: Objective 63x, Zoom 2.3x, Image averaging 2-4x, and airyscan processing.

The percentage of cells showing GFP positivity at the nuclear envelope was counted among mCherry-positive (i.e. competitor expressing) cells in 10 images taken per transfected well.

### Microscopy

Live cells analyzed by microscopy were seeded in transparent Dulbecco’s modified eagle medium without phenol-red (Gibco). Pictures were taken with either a spinning disk confocal microscope (Zeiss) or with an LSM980-multiphoton confocal microscope allowing high-resolution microscopy. Analysis of the image was made using the Zen system image analysis software (Zeiss). Picture of HeLa GFP-NES & RFP-NLS were adjusted using the automatic “best fit”, all other pictures were taken with the same exposure time, image brightness and contrast.

### 3D Antibody production and purification

*Escherichia coli* C41 pLys carrying a pOPTH plasmid coding for a 6xHis-tagged catalytically inactive (GDD->DAA mutant) 3D polymerase was incubated overnight at 37°C in 10ml of tryptic soy broth supplemented with 30 µg/ml of Chloramphenicol and 250 µg/ml of ampicillin (TSB-C-A). This culture was used to inoculate 1L of TSB-C-A and cultured at 37°C. Once OD600 reached 0.8, bacteria were stimulated with 0.5mM of IPTG and cultured for 16h at 20°C. Pellets were resuspended in 150ml of cold resuspension solution (50mM Tris pH 8.4, 150mM NaCl, imidazole 50mM supplemented with 1mM of benzamidine, 1µM pepstatin, 1µg/ml leupeptin, PMSF 1mM). Cells were French press lysed (800 PSI), lysate was then sonicated, centrifuged and the supernatant was cleared with a 0.45 µM filter before being applied on a Nickel affinity column. A HisTrap High Performance column (Cytiva 17-5247-01, Sigma) was rinsed with 2×5ml of H_2_O and 2×5ml of resuspension solution before applying the lysate (1ml/minute). The column was then rinsed with 2×5ml of: resuspension solution R1 (50mM Tris pH8.4, 500mM NaCl, 50mM Imidazole) and R2 (50mM Tris pH8.4, 500mM NaCl). His tagged 3D was eluted with 50mM Tris pH8.4, 500mM NaCl, 300mM Imidazole in 1ml fractions supplemented with 1 volume of 50mM Tris pH8.4, 500mM NaCl, 30% glycerol. After concentration and purity testing, fractions were snap frozen, in aliquots containing 0.1mg of protein.

A classical immunization program (Eurogentec, Belgium) was followed with 2 rabbits (ref AS-PNOR-3MORAB 87-Day program). Briefly, rabbits were immunized at days 0, 14, 28 and 58 with 0.1mg of protein coupled to Freund’s adjuvant. Blood was drawn on days 0 (pre-immune), 66 (small bleed) and 87 (final bleed). Pre-immune, large bleed and final bleed sera were tested by western blot against infected and non-infected cell lysates for specificity.

For purification of a batch of anti-3D antibodies, 10 mg of purified 3D-His protein was dialyzed in 0.1M NaCO_3_ pH 8.4, 0.5M NaCl and coupled to 0.75g of CNBr beads (C9142-5G, Sigma) according to the manufacturer’s instructions. Beads were rinsed with 3×3ml of R solution (50mM Tris pH8, 0.5M NaCl) before applying 2ml of rabbit sera mixed with 1 volume of NaN_3_ 0.2% (w/v), NaCl 0.5M, Tween 20 0.6% (v/v) and filtered with a 0.2µm filter. Non-specific antibodies were removed with 3×3ml of R solution before eluting 3D specific antibodies in 4×1ml fractions with 4M MgCl_2_, 100mM HEPES pH6. Immediately after elution, antibodies were dialyzed in 50mM Tris pH7.4, 150mM NaCl, 0.2% Tween 20 (v/v). After quantification, antibodies were stored in 50% glycerol supplemented with protease free BSA and NaN_3._

## Data availability

All data are contained in the article. Biological material is available upon request.

## Acknowledgements

We are very grateful to Stephane Messe (who sadly passed away in spring 2023) for consistently great technical assistance. We thank Patrick van der Smissen (PICT platform, UCLouvain) for help and training in confocal microscopy.

## Authors’ contribution

Conceptualization: BLP, TM,

Data curation: BLP-MV

Formal analysis: BLP, FW, TM

Funding acquisition: TM

Investigation: BLP, FW, MV, CD, RM, FS, TM

Methodology: FS, MV, RM

Supervision: TM Visualization: BLP, RM

Writing – original draft:BLP, TM, RM.

Writing – review & editing: BLP, FW, MV, CD, RM, FS, TM

FW performed most of the experiments related to Figs 1, 2, 3, 4 & 6. BLP contributed to the conceptualization and analysis of the work, to some constructs and to experiments related to Figs 3 × 7. CD raised a polyclonal anti-3D antibody; FS, MV, and RM were involved in defining, implementing and analyzing AlphaFold predictions. BLP and TM contributed the most to drafting manuscript and figures. All authors contributed to some extent to text edition.

